# Single-Cell Translation and Apoptosis Profiling to Define Human CD34^+^ Cell Response to Specific Factors

**DOI:** 10.64898/2026.05.21.726964

**Authors:** Dan Li, Karin Gustafsson, Jelena Milosevic, Anna Kiem, David T Scadden

**Author notes:** **Correspondence to:** David T Scadden.

## Abstract

Global mRNA translation is a defining functional property of hematopoietic stem cells (HSCs) and is increasingly recognized as a critical axis of dysregulation in myelodysplastic syndromes (MDS) and other clonal hematopoietic disorders. Yet the quantitative measurement of protein synthesis at single-cell resolution across phenotypically defined HSPC subpopulations, in parallel with apoptotic state, is technically challenging. Here we describe and validate a single-tube flow cytometry protocol that simultaneously quantifies global protein synthesis by O-propargyl-puromycin (OP-Puro) incorporation and intracellular cleaved Caspase-3 with cell immunophenotyping across the canonical CD34^+^ HSPC hierarchy in cryopreserved human cord blood (CB) CD34^+^ cells. The protocol enables quantitative assessment of key dynamic cell processes in defined subsets of primary hematopoietic cells on a standard flow cytometer. We apply this assay to a four-condition factor-omission analysis of the canonical SR1 + UM729 + dmPGE2 *ex vivo* expansion cocktail across three independent CB donors. The analysis assigns each compound a distinct functional profile: UM729 constrains protein synthesis and supports apoptotic priming across the hierarchy; SR1 maintains a pro-survival state without modulating translation; and dmPGE2 promotes HSC cycling and progressive exit from the primitive state, with minimal direct effect on the translation or apoptotic axes measured here. This analysis resolves three mechanistically distinct small-molecule signatures using a protocol directly transferable to clinical biobank specimens. We propose it as a functional-state analytic platform that may be useful for patient-derived CD34^+^ cells from MDS and other myeloid neoplasms in which translational dysregulation is a recognized pathological feature.

## 1. INTRODUCTION

Allogeneic hematopoietic stem cell transplantation (HSCT) is curative for hematologic malignancies and select non-malignant disorders, but its broader application is constrained by donor availability. Umbilical cord blood (CB) provides an alternative graft source whose clinical use has historically been limited by the modest stem cell content of a single CB unit relative to adult-recipient cell-dose requirements (Broxmeyer, 2010). *Ex vivo* expansion of CB CD34^+^ hematopoietic stem and progenitor cells (HSPCs) has therefore become a central strategy for extending CB transplantation to adult patients, and three small molecules, the AhR antagonist StemRegenin 1 (SR1; Boitano et al., 2010), the pyrimidoindole UM171 and its analog UM729 (Fares et al., 2014; Chagraoui et al., 2021), and the prostaglandin analog 16,16-dimethyl PGE_2_ (dmPGE2; Hoggatt et al., 2009; Goessling et al., 2011), have together defined the current standard expansion cocktail, with clinical efficacy demonstrated for each component (Wagner et al., 2016; Cohen et al., 2020; Cutler et al., 2013). Despite this clinical maturation, the contribution of each compound to the intracellular state of the expanding HSPC population has not been systematically dissected at single-cell resolution across the canonical HSPC hierarchy.

Among the intracellular axes most informative of HSPC functional state, global mRNA translation is uniquely positioned. Translation is a defining functional property of HSCs: freshly isolated HSCs maintain uniquely low rates of global protein synthesis relative to their downstream progeny, and this translational restraint is essential for self-renewal as both genetic increases (e.g., Pten loss) and decreases (e.g., Rps14 haploinsufficiency) in protein synthesis impair HSC function (Signer et al., 2014; Hidalgo San Jose et al., 2020). Translation captures the integrated output of cytokine and small-molecule signaling through PI3K/AKT/mTOR, JAK/STAT, MAPK and Wnt convergence on the eIF4F complex and ribosome biogenesis and is mechanistically coupled to lineage commitment through ribosome-content–dependent translation programs (Khajuria et al., 2018).

Critically, translational dysregulation is a recurrent and increasingly recognized theme in clonal hematopoietic disorders, including Diamond-Blackfan anemia, Shwachman-Diamond syndrome, 5q− MDS, and TET2-mutant clonal hematopoiesis and MDS, where altered protein synthesis programs are emerging as both biomarkers and candidate therapeutic vulnerabilities (Saba et al., 2021; Fabbri et al., 2021). A robust per-cell translation readout in phenotypically defined human HSPCs is therefore of substantial value not only for expansion biology but also for the dissection of disease-relevant translational programs in patient-derived material.

Apoptotic priming is the second axis we measure in parallel, for three reasons. First, apoptotic loss is the primary mode of HSPC depletion during *ex vivo* culture and the specific bottleneck that the dmPGE2 component of the cocktail aims to address. Second, apoptotic priming is mechanistically coupled to translational regulation: cap-dependent global translation is suppressed during apoptotic execution while IRES-mediated translation of selective pro-survival and pro-apoptotic mRNAs persists (Holcik and Sonenberg, 2005), such that the joint measurement of translation and apoptosis provides a more informative readout of cellular state than either alone. Third, cleaved Caspase-3 (cCasp3) detection by flow cytometry captures cells that have committed to apoptotic execution and is a quantitative single-cell readout of apoptotic priming, with the BCL-2 family balance that determines priming representing a major therapeutic target in hematologic malignancies including MDS and AML (Bhatt et al., 2020).

Here, we describe and validate a seven-step, single-tube flow cytometric protocol that simultaneously measures global translation, multi-parameter HSPC immunophenotype, viability, and intracellular cCasp3 in human CB CD34^+^ cells. The protocol permits analyses not accessible by bulk or surface-only methods. It integrates OP-Puro labeling for global translation rate (Liu et al., 2012; Forester et al., 2018), surface staining of the canonical HSPC hierarchy with EPCR refinement of the most primitive HSC fraction (Fares et al., 2017; Anjos-Afonso et al., 2022), fixation/permeabilization, copper-catalyzed click chemistry for OP-Puro detection, and intracellular antibody staining within a single workflow. We apply the assay to a four-condition factor-omission analysis of the SR1 + UM729 + dmPGE2 cocktail across three independent CB donors, demonstrating that the assay resolves three mechanistically distinct small-molecule signatures. The protocol is fully compatible with cryopreserved primary human CD34^+^ cells and is readily adaptable to other intracellular protein targets, including transcription factors, signaling molecules, and disease-relevant epigenetic regulators, by substitution of the intracellular antibody, providing a methodological framework for the joint dissection of translation and apoptosis in clinical biobank specimens from patients with MDS and related myeloid neoplasms.

## 2. RESULTS

### 2.1 A seven-step single-tube protocol enables simultaneous quantification of translation, immunophenotype, viability, and apoptosis in primary human CD34^+^ cells

We developed a seven-step, single-tube flow cytometric protocol (**Figure 1A**) for simultaneous quantification of global translation rate, multi-parameter HSPC immunophenotype, cell viability, and intracellular protein expression in human CB CD34^+^ cells. The workflow sequentially incorporates: (Step 1) OP-Puro labeling of nascent polypeptides at 20 µM for 30 min at 37 °C in complete culture medium; (Step 2) surface antibody staining for an HSPC-resolving panel (CD34, CD38, CD90, CD45RA, EPCR); (Step 3) labeling with a fixable amine-reactive viability dye to discriminate live and dead cells through subsequent fixation; (Step 4) fixation in paraformaldehyde and permeabilization in saponin-based buffer; (Step 5) copper-catalyzed azide–alkyne click reaction conjugating a fluorophore-azide to the OP-Puro–labeled nascent peptides; (Step 6) intracellular antibody staining (in this study, anti-cleaved Caspase-3, with a parallel IgG isotype-stained tube as a matched control); and (Step 7) flow cytometric acquisition on a standard analytical cytometer.

**Figure 1.**
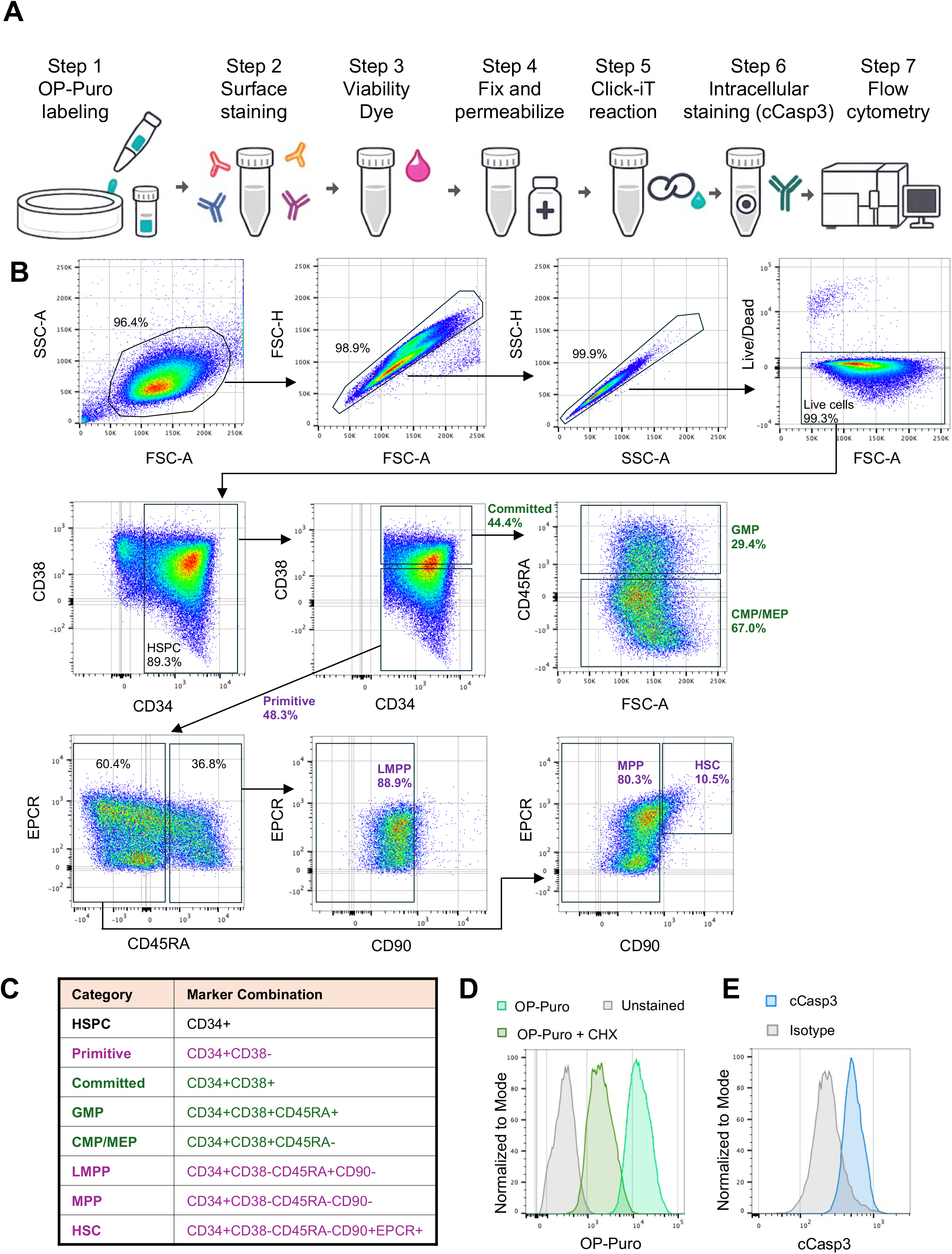
Single-tube flow cytometric protocol for simultaneous quantification of global translation, HSPC immunophenotype, viability, and intracellular protein in primary human CD34^+^ cells. (A) Seven-step workflow. (B) Hierarchical gating strategy resolving the canonical HSPC hierarchy with CD201/EPCR refinement of the HSC fraction. (C) Surface marker combinations defining each population. (D) Representative OP-Puro histogram with cycloheximide (CHX) and unstained controls confirming translation-dependent OP-Puro signal. (E) Representative cCasp3 histogram with matched IgG isotype control.

The hierarchical gating strategy (**Figure 1B and 1C**) resolves the canonical CB HSPC hierarchy at high resolution: from scatter-defined intact cells through singlets and live cells, to total CD34^+^ HSPCs and primitive CD34^+^CD38^−^ versus committed CD34^+^CD38^+^ compartments. Within the primitive compartment, CD45RA, CD90 and EPCR discriminate lymphoid-primed multipotent progenitors (LMPPs; CD45RA^+^CD90^−^), multipotent progenitors (MPPs; CD45RA^−^CD90^−^), and HSC (refined by EPCR to identify the most primitive subset; CD34^+^CD38^−^ CD45RA^−^CD90^+^ CD201^+^), enriched for long-term repopulating activity in xenotransplantation assays (Fares et al., 2017; Anjos-Afonso et al., 2022). Within the committed compartment, CD45RA distinguishes granulocyte-monocyte progenitors (GMPs; CD45RA^+^) from common myeloid and megakaryocyte-erythroid progenitors (CMP/MEPs; CD45RA^−^). The single-tube design minimizes inter-sample technical variability and enables direct per-cell correlation of translational and intracellular readouts within each phenotypically defined population. The OP-Puro signal is specifically translation-dependent, abolished by cycloheximide pre-treatment (**Figure 1D**), and the cCasp3 signal is markedly above matched IgG isotype (**Figure 1E**). The protocol is fully compatible with cryopreserved primary human CD34^+^ cells, the standard form in which CB units and patient CD34^+^ samples from clinical biobanks are stored and is readily adaptable to other intracellular protein targets through simple substitution of the Step 6 antibody.

To validate the assay’s sensitivity in detecting biologically meaningful intracellular state changes, we applied it to a four-condition factor-omission analysis of the canonical SR1 + UM729 + dmPGE2 *ex vivo* expansion cocktail. Cryopreserved CB CD34^+^ cells from three independent donors were thawed and cultured for 7 days in StemSpan SFEM supplemented with SCF, FLT3L, TPO, and IL-6 under hypoxic conditions (5% O_2_), with the complete cocktail (“Full”: SR1 1 µM + UM729 35 nM + dmPGE2 1 µM) or one factor omitted (−SR1, −UM729, −dmPGE2; **Table 1**). A parallel kinetic time-course experiment using a focused immunophenotyping panel (without CD38, OP-Puro, or cCasp3 staining) characterized the temporal dynamics of total live cell expansion, CD34^+^CD90^+^ progenitors, and EPCR^+^ HSCs at days 2, 4, 5, and 6 (**Supplementary Figure S1**); these data established that the principal between-condition phenotypic separations emerge by day 4 and are stable through day 6, and they motivated the day-7 endpoint analysis described below.

**Table 1.** Experimental culture conditions.

| Condition | SR1 (1 μM) | UM729 (35 nM) | dmPGE2 (1 μM) | Designation |
| --- | --- | --- | --- | --- |
| Full cocktail | + | + | + | Full |
| Minus SR1 | – | + | + | –SR1 |
| Minus UM729 | + | – | + | –UM729 |
| Minus dmPGE2 | + | + | – | –dmPGE2 |

### 2.2 Endpoint phenotyping reveals stage-selective frequency effects across the HSPC hierarchy

At the day-7 endpoint, cells from the three independent CB donors cultured under each of the four conditions were processed through the full single-tube assay (**Figure 1A**). The expanded antibody panel including CD38 resolved the canonical HSPC hierarchy at fine resolution, separating primitive (CD34^+^CD38^−^) from committed (CD34^+^CD38^+^) compartments and resolving HSC, MPP, LMPP, CMP/MEP, and GMP populations (**Figure 2**).

**Figure 2.**
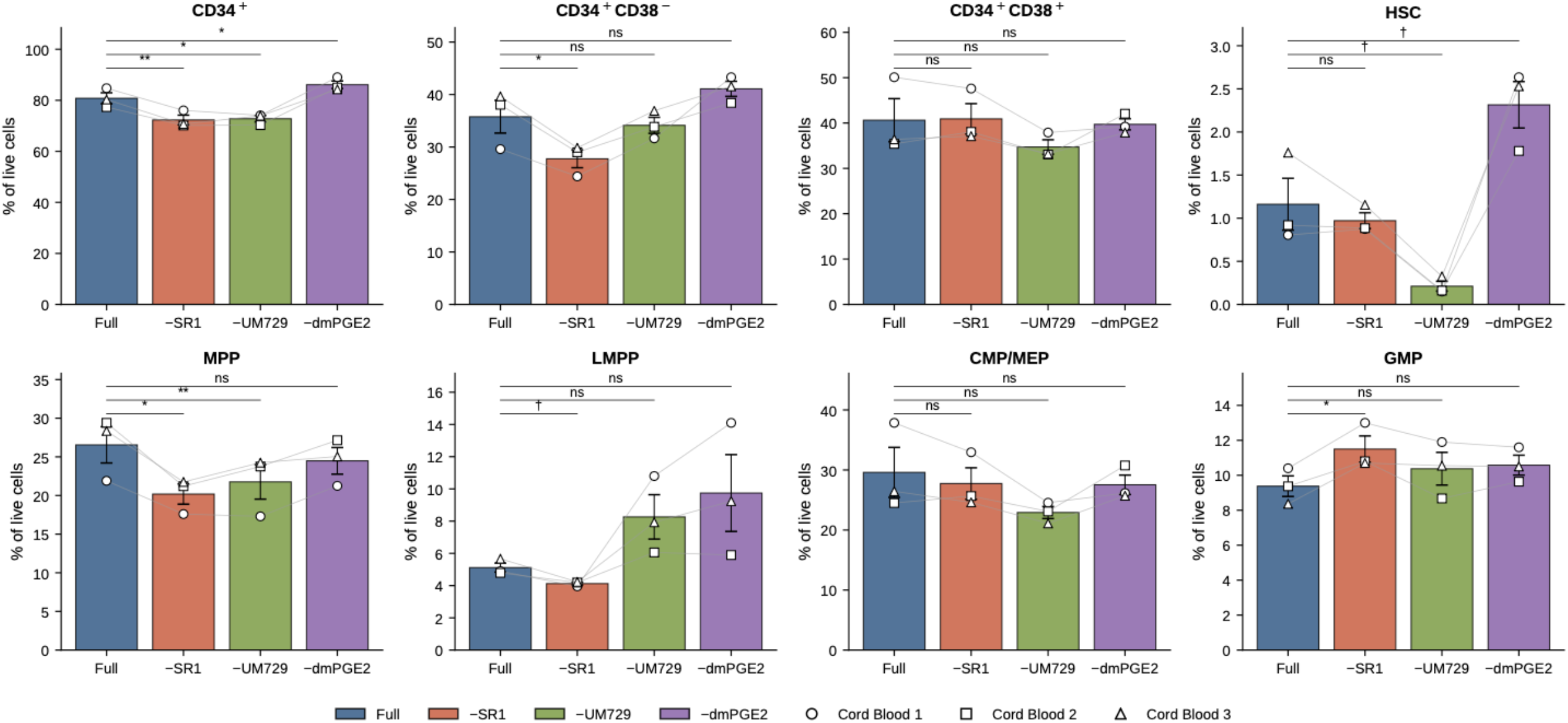
HSPC subpopulation frequencies at day 7 across four cocktail conditions. Population frequencies are expressed as percent of live cells in three independent CB donors. Bars show mean ± SEM (n = 3); individual donors are shown as connected markers (○ Cord Blood 1, □ Cord Blood 2, △ Cord Blood 3). †p < 0.10, *p < 0.05, **p < 0.01, ***p < 0.001 (paired t-test vs. Full cocktail).

The most pronounced compositional effect was on the HSC fraction. UM729 omission reduced this population approximately 5.6-fold (1.16 → 0.21% of live cells; p < 0.001), confirming that UM729 is required for maintenance of the most primitive HSC compartment throughout the culture. Conversely, dmPGE2 omission was associated with an approximately 2-fold increase in EPCR^+^ HSC frequency (1.16 → 2.31%), consistent with the progressive HSC preservation also observed in the kinetic experiment (**Supplementary Figure S1**). The total CD34^+^ frequency was significantly reduced by SR1 omission (80.8 → 72.3%; p = 0.007) and UM729 omission (80.8 → 72.8%; p = 0.027) and trended upward upon dmPGE2 omission (80.8 → 86.1%; p = 0.05).

SR1 omission selectively contracted the most primitive compartments, CD34^+^CD38^−^ primitive cells (35.8 → 27.8%; p = 0.04), MPPs (26.6 → 20.2%; p = 0.04), and LMPPs (5.1 → 4.1%; p = 0.04), while concurrently expanding the GMP fraction (9.4 → 11.5%; p = 0.04), consistent with AhR-driven myeloid commitment in the absence of SR1-mediated AhR antagonism. CMP/MEP frequencies and total CD34^+^CD38^+^ committed-progenitor frequencies were not significantly affected by any single-factor omission.

Having established the compositional landscape of each condition, we asked whether the surviving cells in each condition are translationally and apoptotically equivalent or differ in their intracellular state. Surface phenotyping alone cannot answer this question; we therefore turned to the OP-Puro and intracellular cCasp3 components of the assay.

### 2.3 UM729 restrains global translation across the HSPC hierarchy; SR1 and dmPGE2 do not

Global translation rate was quantified as OP-Puro geometric MFI within each gated population and expressed as fold-change relative to the Full cocktail within each donor and gate (**Figure 3**). Across nearly every population, UM729 omission produced the strongest and most statistically robust translation effect, elevating OP-Puro signal by 16% in bulk CD34^+^ (p = 0.02), 24% in CD34^+^CD38^−^ primitive cells (p = 0.008), 31% in MPPs (p = 0.005), 14% in CMP/MEPs (p = 0.02), and 27% in HSCs (p = 0.06). In contrast, omission of SR1 produced essentially no shift in OP-Puro signal in any population. Omission of dmPGE2 produced small but reproducible OP-Puro elevations in LMPPs (+12%, p = 0.045) with smaller trends elsewhere.

**Figure 3.**
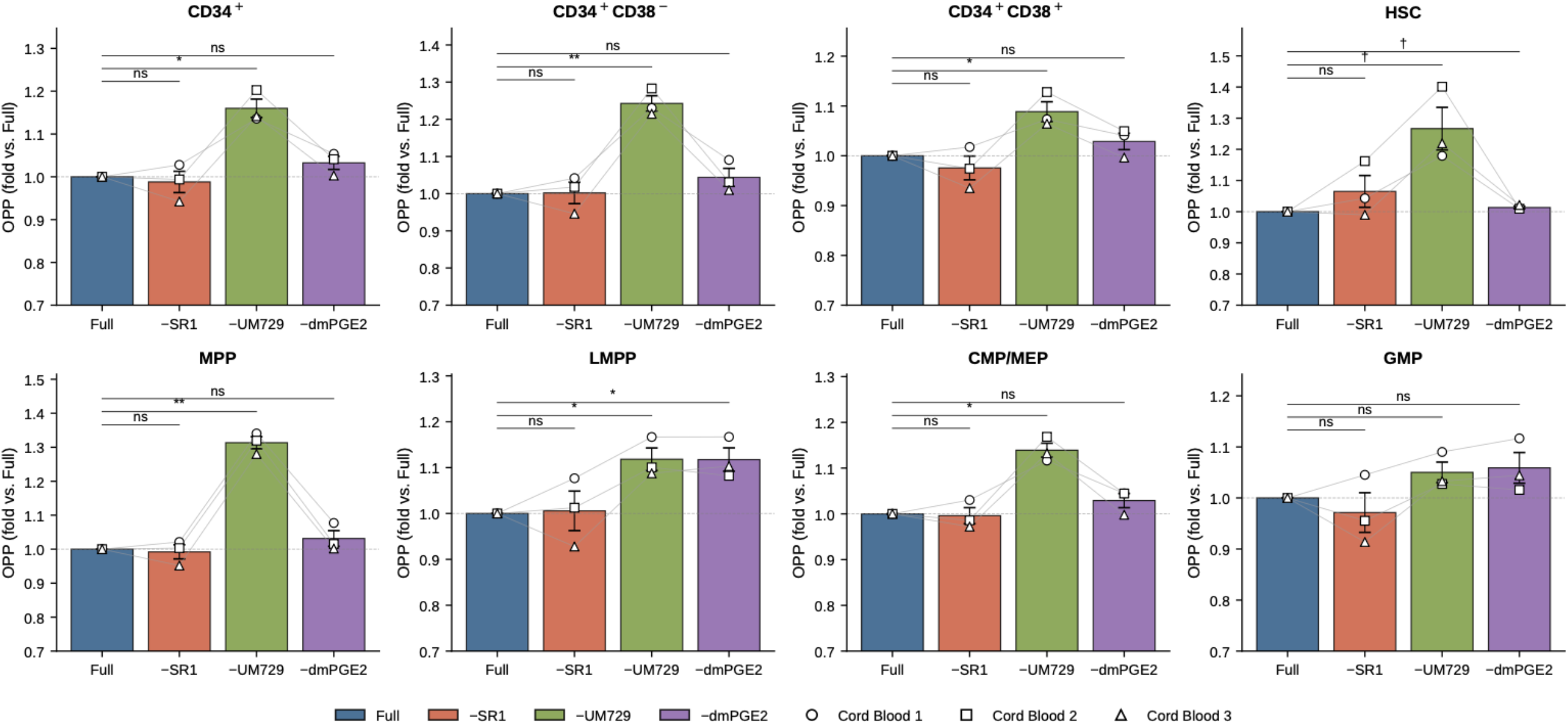
Global translation rate response to factor omission at day 7, presented as OP-Puro geometric MFI fold-change relative to the Full cocktail within each donor and gate. UM729 omission produces the strongest and most consistent elevation of OP-Puro signal across the hierarchy; SR1 and dmPGE2 have minimal effect. Dashed line at fold = 1.0. †p < 0.10, *p < 0.05, **p < 0.01.

These data identify UM729 as the dominant translational modulator in the cocktail. The increase in OP-Puro signal seen with UM729 removal is observed across the entire CD34^+^ hierarchy and is most pronounced in the primitive (CD34^+^CD38^−^) and MPP populations. Together with the simultaneous loss of EPCR^+^ HSCs under the same condition, this suggests that UM729 normally constrains protein synthesis, with the consequent low biosynthetic state linked to maintaining HSC identity.

### 2.4 SR1 acts as a pan-hierarchy survival factor without affecting translation

Apoptotic priming was quantified as the cleaved Caspase-3/IgG geometric MFI ratio and expressed as fold-change relative to Full within each donor and gate (**Figure 4**). The dominant pattern was that SR1 omission elevated cCasp3 ratios across all eight examined populations by 2–10%, with the largest trends in MPPs (+10%), CD34^+^CD38^−^ primitive cells (+9%), and EPCR^+^ HSCs (+9%). While individual population effects did not reach formal statistical significance at n = 3, the directionally consistent elevation across all eight populations is highly suggestive of a generalized pro-survival role for SR1. AhR antagonism by SR1 has been shown to restrict differentiation, likely contributing to lower apoptotic priming as terminal differentiation is limited (Boitano et al., 2010).

**Figure 4.**
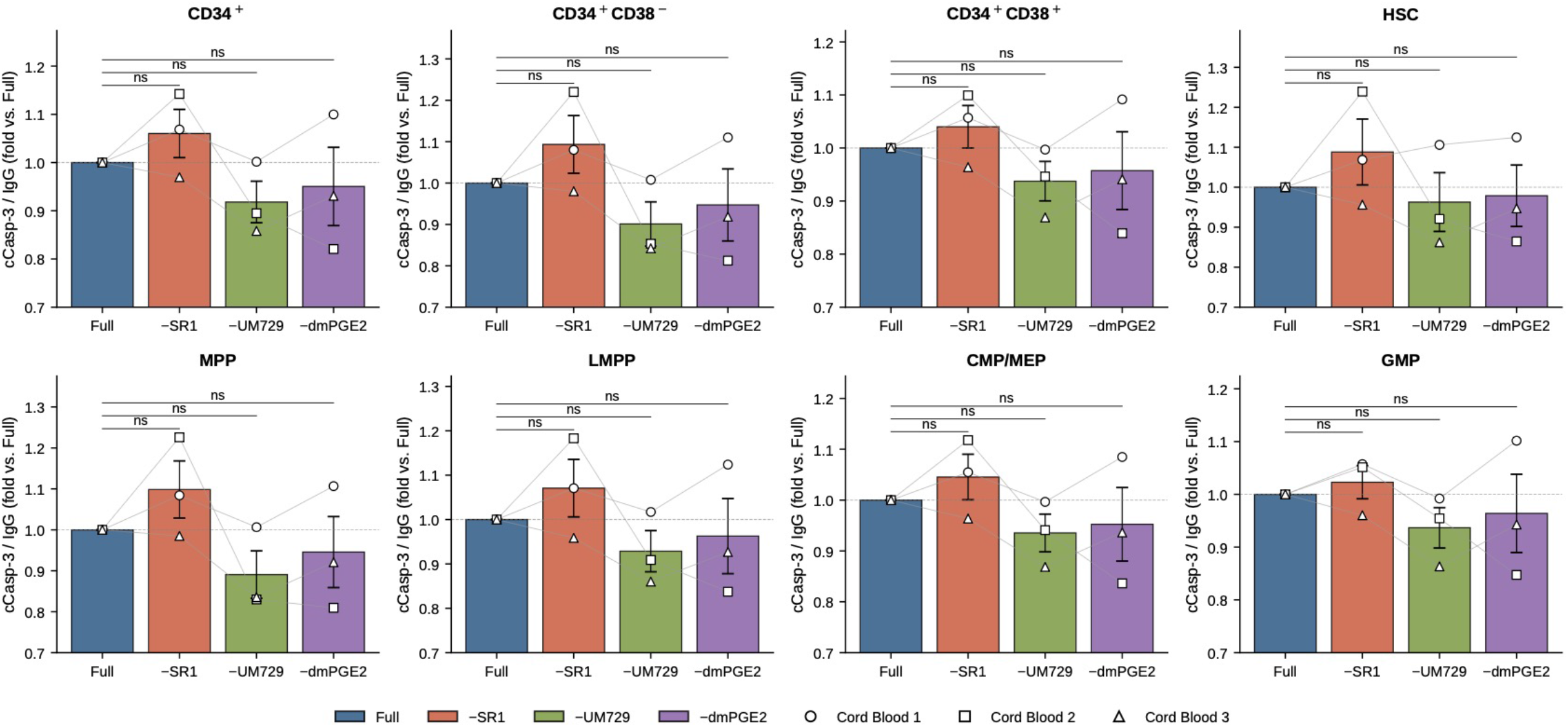
Apoptotic priming response to factor omission at day 7, presented as cleaved Caspase-3 / IgG MFI ratio fold-change relative to the Full cocktail within each donor and gate. SR1 omission produces a directionally consistent elevation across all eight populations. Dashed line at fold = 1.0. †p < 0.10, *p < 0.05.

UM729 omission, by contrast, consistently reduced cCasp3 ratios (−4 to −11% across populations), likely reflecting selective attrition of the most apoptosis-primed primitive cells over the 7-day culture in the absence of UM729-driven self-renewal support, consistent with the parallel collapse of the EPCR^+^ HSC compartment (**Figure 2**) under this condition.

dmPGE2 omission produced small reductions in cCasp3 (−2 to −5% across populations). This minimal apoptotic effect is consistent with the dominant chronic action of dmPGE2 on HSPCs operating through EP2/EP4 → cAMP → CREB-mediated chromatin remodeling (Fast et al., 2015; Sporrij et al., 2023) and Wnt/β-catenin signaling (Goessling et al., 2009) rather than the apoptotic axis; the previously characterized acute anti-apoptotic effect of PGE2 via Survivin induction (Hoggatt et al., 2009) is most evident with brief (1–2 hour) pulse exposure rather than the chronic 7-day exposure used here.

Together, the translation and apoptosis readouts assign each cocktail component a distinct functional profile. The resolution of three compound-specific profiles validates the assay’s ability to distinguish distinct intracellular effects with the relative ease and precision of a flow cytometric assay.

## 3. DISCUSSION

### 3.1 A single-tube assay for joint translation and apoptosis profiling of human HSPCs

The seven-step protocol described here enables simultaneous single-cell quantification of global protein synthesis, multi-parameter HSPC immunophenotype, cell viability, and apoptotic priming in human CB CD34^+^ cells, using a standard analytical flow cytometer and commercially available reagents. The integration of OP-Puro click chemistry with surface immunophenotyping and intracellular cCasp3 staining in a single tube minimizes inter-sample technical variability and enables direct correlation of translational and apoptotic state within phenotypically defined populations. The within-gate fold-change framework adopted throughout normalizes each gate’s response to factor omission against that gate’s own full cocktail value within each donor, controlling for inter-donor variability. We deliberately avoid cross-population comparisons of absolute OP-Puro and cCasp3 magnitudes because, after 7 days of *ex vivo* culture, each immunophenotypically defined population has diverged substantially from its fresh *in vivo* counterpart along a population-specific adaptation trajectory driven by the cytokine combination, the small-molecule cocktail, and the culture environment itself; the cultured “HSC”, “MPP”, and “GMP” gates therefore reflect the convergent result of intrinsic population identity and 7 days of differential culture adaptation. The within-gate fold-change against the Full reference cleanly isolates the experimental variable of interest, cocktail composition, from this culture-adaptation confound.

### 3.2 Two findings warranting further mechanistic evaluation

Of the three patterns we observed (Sections 2.3–2.4), two have implications worth further discussion. The first concerns UM729, which appears to act as a translational restraint: removing it increased OP-Puro signal across the hierarchy while at the same time depleting the EPCR^+^ HSC compartment. This suggests that UM729 holds back the biosynthetic activation that the cytokine cocktail (SCF + FLT3L + TPO + IL-6) would otherwise drive, even while it promotes HSC self-renewal through KBTBD4–CRL3-mediated proteolysis of LSD1/CoREST (Chagraoui et al., 2021). To our knowledge, this pairing of translational restraint and self-renewal has not been described for UM171/UM729-class compounds and merits direct mechanistic follow-up, particularly given the central role of ribosome biogenesis in HSC differentiation (Khajuria et al., 2018).

Second, the dmPGE2 result invites reframing relative to the acute-pulse literature. dmPGE2 was developed and clinically validated as a brief (1–2 hour) pre-transplant pulse that enhances HSC homing and survival (Hoggatt et al., 2009; Cutler et al., 2013); in our 7-day protocol it was instead present continuously at 1 µM. Under this chronic regimen, dmPGE2 omission was associated with an approximately two-fold increase in EPCR^+^ HSC frequency that, while directionally consistent across the three donors, did not achieve statistical significance. One possible interpretation is that chronic EP2/EP4 → cAMP/PKA and Wnt/β-catenin signaling, distinct from the acute pulse context, drives HSC cycling and progressive exit from the most primitive state. Functional confirmation by limiting-dilution xenotransplantation will be required to distinguish true preservation of repopulating activity from phenotypic retention; if validated, the finding would motivate reconsidering whether dmPGE2 is best deployed as a brief pre-transplant supplement rather than a chronic culture additive.

### 3.3 Application to MDS and other clonal hematopoietic disorders

The principal motivation for this assay framework is its direct applicability to CD34^+^ cells from patients with clonal hematopoietic disorders in which translational dysregulation is a recurrent pathological feature. In MDS, ribosome biogenesis defects (5q− syndrome with RPS14 haploinsufficiency) and germline ribosomal protein mutations (Diamond-Blackfan anemia, Shwachman-Diamond syndrome) drive translational programs contributing to ineffective hematopoiesis and clonal evolution (Khajuria et al., 2018; Saba et al., 2021), and emerging evidence implicates additional recurrent MDS-associated regulators including TET2 in translational control (Zou et al., 2024). The seven-step protocol is directly transferable to the cryopreserved CD34^+^ patient samples archived in hematologic malignancy biobanks, and substitution of the intracellular Step 6 antibody enables quantification of any intracellular protein of interest, including BCL-2 family members, ribosomal proteins, and translation initiation factors, in parallel with the translational and apoptotic readouts. The framework therefore positions the joint single-cell measurement of translation and apoptosis as a practical readout for the dissection of disease-relevant intracellular programs in patient-derived MDS material.

### 3.4 Limitations and future directions

There are two additional limitations we acknowledge in this study, beyond the culture-adaptation caveat discussed in Section 3.1. First, immunophenotypic populations were not validated functionally by xenotransplantation in this study, and the interpretation depends on prior work establishing these surface markers as predictive of repopulating activity. Second, CD90 is a GPI-anchored protein partially vulnerable to detergent permeabilization, potentially introducing systematic signal loss in CD90^+^ gates; CD201/EPCR refinement partially mitigates this.

Future work will include systematic dose–response analyses, temporal mapping of the translational and apoptotic readouts beyond the day-7 endpoint, and, most importantly, extension to patient-derived HSPCs from MDS and related myeloid neoplasms.

## 4. METHODS

### 4.1 Cord blood CD34^+^ cells

Cryopreserved purified human cord blood CD34^+^ cells from three independent donors were obtained from MGB (IRB Protocol#: 2010P002371). Cells were thawed in a 37 °C water bath, slowly diluted into pre-warmed StemSpan SFEM medium, centrifuged at 300 × g for 8 min, and resuspended in pre-warmed expansion medium for immediate seeding. CB collection and use were approved by MGH (IRB Protocol#: 2010P002371).

### 4.2 Expansion culture

CD34^+^ cells were seeded at 1 × 10^5^ cells/mL in StemSpan SFEM medium (StemCell Technologies, #09600) supplemented with recombinant human cytokines (all Peprotech): SCF (#300-07; 100 ng/mL), FLT3L (#300-19; 100 ng/mL), TPO (#300-18; 20 ng/mL), and IL-6 (#200-06; 20 ng/mL). Penicillin/streptomycin (Gibco, #15140-122; 1× working concentration) and L-glutamine (Gibco, #25030-081; 1× working concentration) were included throughout. The complete (“Full”) small-molecule cocktail contained StemRegenin 1 (SR1; Cayman Chemical or equivalent; 1 µM), UM729 (StemCell Technologies or equivalent; 35 nM), and 16,16-dimethyl prostaglandin E2 (dmPGE2; Cayman Chemical, #14750; 1 µM). The three additional conditions were established by omitting one of the three compounds (**Table 1**). Cultures were maintained at 37 °C, 5% CO_2_, and 5% O_2_ in a humidified hypoxia incubator for 7 days, with media replaced every other day. n = 3 independent CB donors, each tested across all four conditions.

### 4.3 Kinetic time-course experiment

To characterize the temporal dynamics of HSPC expansion under each cocktail condition, an independent time-course experiment was performed using a focused immunophenotyping panel. Cord blood CD34^+^ cells were thawed and seeded as described above under the four conditions described in Section 4.2 and **Table 1**. Aliquots were harvested for flow cytometric analysis on days 2, 4, 5, and 6. Surface staining was performed with antibodies against CD34, CD90, CD45RA, and EPCR; the CD38, OP-Puro, and intracellular cCasp3 components of the full assay were not included in this focused panel. Live cells were identified by viability-dye exclusion as in Section 4.4. The primitive HSC fraction was defined as CD34^+^CD45RA^−^CD90^+^EPCR^+^, which differs from the main experiment HSC gate by omission of the CD38^−^ criterion; both definitions identify EPCR-enriched primitive HSC populations consistent with Fares et al. (2017).

### 4.4 OP-Puro labeling and seven-step flow cytometric protocol

The OP-Puro labeling and click-chemistry detection steps adapt the *in vivo* mouse flow cytometric protocol for cell-type-specific quantification of protein synthesis (Hidalgo San Jose and Signer, 2019), here modified for *ex vivo* culture of human cord blood CD34^+^ cells and integrated with multi-color HSPC immunophenotyping and intracellular cleaved Caspase-3 staining within a single tube.

On day 7, cells were collected, counted, and processed through the seven-step protocol (**Figure 1A**). **Step 1 (OP-Puro labeling):** cells were incubated with O-propargyl-puromycin (20 µM) for 30 min at 37 °C in expansion medium. A parallel cycloheximide pre-treatment control (100 µg/mL, 15 min before OP-Puro addition) was included to define translation-independent background. **Step 2 (surface staining):** cells were washed in PBS + 2% FBS and stained on ice for 30 min with the following antibody panel: anti-CD34-APC-Cy7 (Clone: 581; BioLegend Catalog: 343514), anti-CD38-PE-Cy7 (Clone: HIT2; BioLegend Catalog: 303516), anti-CD90-BV711 (Clone: 5E10; BioLegend Catalog: 328139), anti-CD45RA-BUV737 (Clone: HI100; BD Biosciences Catalog: 564442), and anti-CD201-APC (Clone: RCR-401; BioLegend Catalog: 351906). **Step 3 (viability):** cells were washed and incubated with LIVE/DEAD Fixable [Blue] Dead Cell Stain (Thermo Fisher; UV-excited, emission 450 nm; Catalog: L34962) in PBS for 20 min at room temperature (RT) protected from light. **Step 4 (fixation/permeabilization):** cells were fixed in 4% methanol-free formaldehyde for 10 min at RT in dark and permeabilized in 0.1% saponin / 3% FBS in PBS for 10 min at RT. **Step 5 (click reaction):** the Click-iT Plus reaction cocktail (containing AlexaFluor-azide 405, CuSO_4_) was applied for 30 min at RT protected from light, then washed twice in permeabilization buffer. **Step 6 (intracellular staining):** cells were stained with anti-cleaved Caspase-3-AlexaFluor 488 (R&D systems Catalog: IC835G) or matched IgG isotype-AF488 control for 30 min at RT in permeabilization buffer, then washed three times in permeabilization buffer. **Step 7 (acquisition):** samples were acquired on a FACS Aria flow cytometer with appropriate laser and filter configurations. A minimum of 100,000 live-cell events were collected per sample. Compensation was performed using UltraComp eBeads (eBioscience) and matrix calculated in FlowJo v10 (BD Biosciences).

### 4.5 Flow cytometry data analysis

Data were analyzed in FlowJo v10. The gating hierarchy was: scatter (FSC-A vs. SSC-A) → singlet discrimination (FSC-H vs. FSC-A, SSC-H vs. SSC-A) → viability gate (LIVE/DEAD-negative) → CD34^+^ → CD38 split (CD34^+^CD38^−^ vs. CD34^+^CD38^+^) → CD90/CD45RA split within each, with CD201 refinement of the CD90^+^CD45RA^−^ HSC gate (**Figure 1B**). Geometric mean fluorescence intensity (MFI) was extracted for OP-Puro (BV421/405-azide channel), cCasp3 (AF488 channel), and FSC-A in each gate. cCasp3-specific signal was quantified as the ratio of cCasp3 geometric MFI in the cCasp3-stained tube to that in the matched IgG-stained tube. OP-Puro signal per gate was computed as the mean of OP-Puro geometric MFI values from the cCasp3 and IgG tubes (which should not differ by experimental design). For each metric, fold-change relative to the Full cocktail was computed within each donor and gate. Population frequencies were expressed as percent of live cells.

### 4.6 Statistical analysis

All statistical comparisons were performed in Python (pandas, scipy) and confirmed in GraphPad Prism v10. For the day-7 endpoint experiment, paired t-tests were used to compare each factor-omission condition to the Full cocktail within matched donors; two-sided p-values are reported. For the kinetic experiment (**Supplementary Figure S1**), one-way ANOVA was performed at each time point followed by Dunnett’s multiple-comparison test against the Full cocktail. Significance thresholds: †p < 0.10, *p < 0.05, **p < 0.01, ***p < 0.001, ****p < 0.0001. Given the sample size of n = 3 in the endpoint experiment, marginal p-values (0.05 < p < 0.10) are reported and interpreted with reference to directional consistency across donors. Data are presented as mean ± standard error of the mean (SEM).

## AUTHOR CONTRIBUTIONS

D.L.: Conceptualization, Data curation, Formal analysis, Investigation, Methodology, Visualization, Writing – original draft. K.G., J.M., A.K.: Investigation, Methodology, Writing – review & editing. D.T.S.: Conceptualization, Funding acquisition, Project administration, Supervision, Writing – review & editing.

## ACKNOWLEDGMENTS

We were supported with expert technical assistance by the HSCI-CRM Flow Cytometry facility. D.T.S. was supported by the Gerald and Darlene Jordan Professor Chair, the Harvard Stem Cell Institute, and PO1 HL131477, PO1 HL183483 and U19 HL156247. Cord blood units were obtained through IRB Protocol#: 2010P002371.

## COMPETING INTERESTS

D.T.S. is a director of Lightning Biotherapeutics, Agios Pharmaceuticals and Editas Medicine; is a founder and stockholder of Lightning Biotherapeutics and Fate Therapeutics; and is a consultant for Moderna and GV. The remaining authors declare no competing financial interests.

## DATA AVAILABILITY

All flow cytometric data (FCS files and FlowJo workspaces) and processed source data underlying the figures of this study are available from the corresponding author upon reasonable request.

## SUPPLEMENTARY FIGURE LEGENDS

**Supplementary Figure S1.**
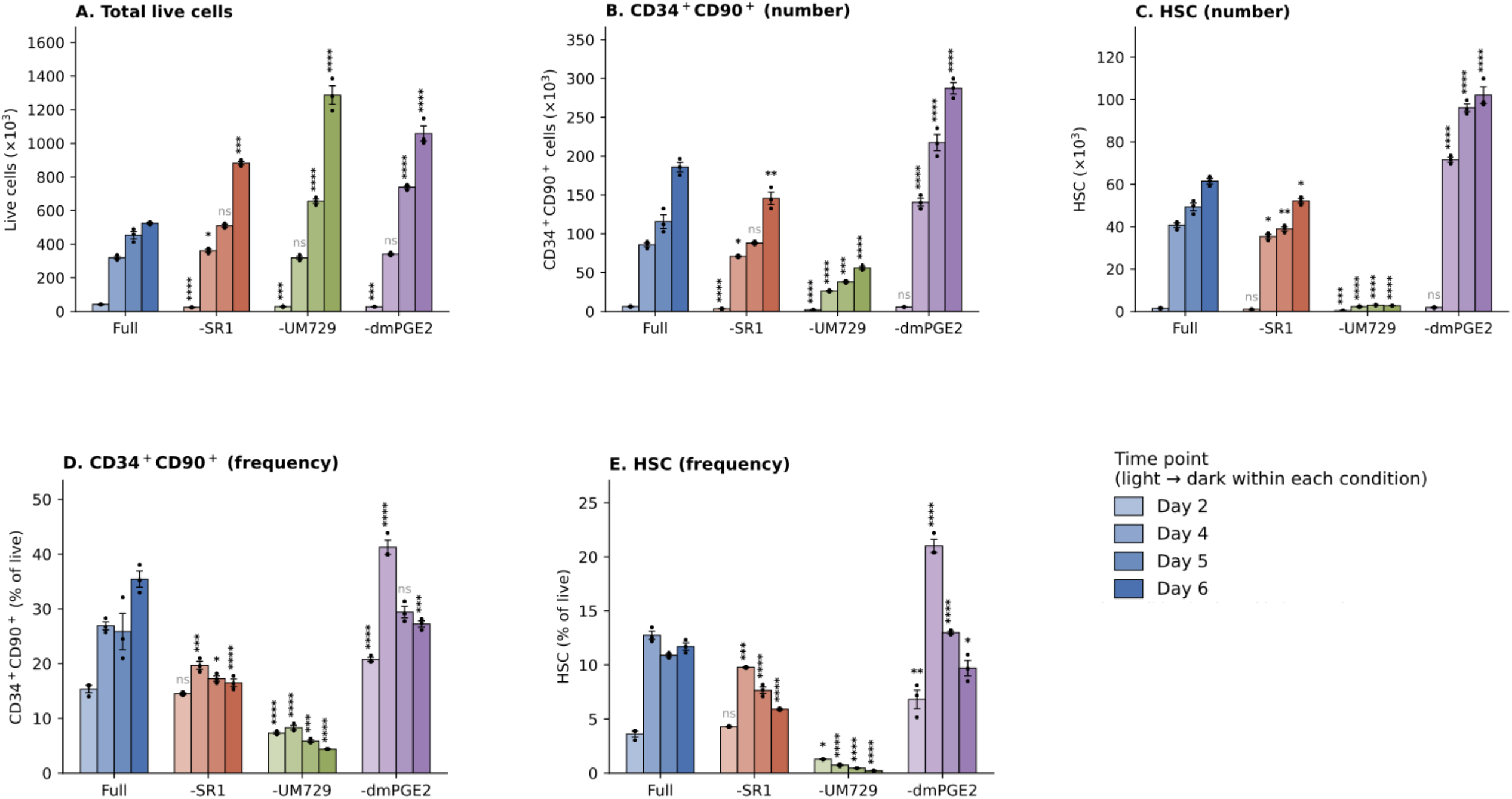
Kinetic analysis of HSPC expansion under four cocktail conditions. (A) Total live cell count, (B) absolute CD34^+^CD90^+^ progenitor count, (C) absolute HSC (CD34^+^CD90^+^CD45RA^−^EPCR^+^) count, (D) CD34^+^CD90^+^ frequency (% of live cells), and (E) HSC frequency (% of live cells) at days 2, 4, 5, and 6 under each of the four cocktail conditions. Within each condition, bars are shaded light → dark for days 2 → 6; each condition is plotted in its own hue (Full = blue, −SR1 = coral, −UM729 = olive, −dmPGE2 = purple) matching the color scheme of Figures 2–4. Bars show mean ± SEM with individual replicates (n = 3) overlaid as dots. Statistics: one-way ANOVA at each time point followed by Dunnett’s multiple-comparison test against the Full cocktail. *p < 0.05, **p < 0.01, ***p < 0.001, ****p < 0.0001; ns, not significant. This experiment used a focused immunophenotyping panel (CD34, CD90, CD45RA, EPCR/CD201) without CD38, OP-Puro, or cCasp3 staining; the HSC gate is therefore defined as CD34^+^CD90^+^CD45RA^−^EPCR^+^ (omitting the CD38^−^ criterion used in the day-7 endpoint experiment).

